# The histone methyltransferase SETD2 regulates HIV expression and latency through a post-transcriptional mechanism

**DOI:** 10.1101/2023.12.15.571871

**Authors:** Cameron Bussey-Sutton, Airlie Ward, Anne-Marie W Turner, Jackson J Peterson, Ann Emery, Arturo SR Longoria, David M Margolis, Brian D Strahl, Edward P Browne

## Abstract

HIV can enter a state of transcriptional latency in CD4 T cells, allowing the virus to evade the host immune system and persist during antiretroviral therapy. Thus, understanding the mechanisms that drive HIV latency, and developing strategies to reactivate viral expression in latently infected cells, are key goals for achieving a cure for HIV. The full spectrum of mechanisms behind the regulation of HIV expression and latency are unclear but include covalent modifications to cellular histones that are associated with the integrated provirus. Here we investigate the role of the SETD2 histone methyltransferase, which deposits H3K36 trimethylation (H3K36me3) cotranscriptionally at genes, in HIV infection. We show that prevention of H3K36me3 through addition of a potent and selective inhibitor of SETD2 (EPZ-719) in human T cells leads to reduced post-integration viral gene expression and accelerated the emergence of a latently infected pool within a population of infected cells. CRISPR/Cas9-mediated knockout of SETD2 in HIV infected primary CD4 T cells confirmed the role of SETD2 in HIV expression. Intriguingly, EPZ-719 exposure also enhanced responsiveness of latently infected cells to latency reversal with the HDAC inhibitor vorinostat. Transcriptomic profiling of EPZ-719 exposed HIV-infected cells identified numerous cellular pathways impacted by EPZ-719. Finally, we show SETD2 inhibition does not affect HIV viral transcription, but instead leads to a shift in the pattern of viral RNA splicing – a result that would be predicted to reduce HIV expression. These results identify SETD2 and H3K36me3 as novel regulators of HIV expression and latency through a post-transcriptional mechanism.

## Introduction

Antiretroviral therapy (ART) targeting of key HIV enzymes has been enormously successful at treating HIV infection and has allowed people with HIV (PWH) to lead relatively normal lives. Nevertheless, a cure for HIV has remained elusive, and interruption of ART results in rapid viral rebound [1,2]. The details of how HIV is able to persist during decades of therapy are not entirely clear, but likely involves the ability of HIV to enter a state of latent infection in memory CD4 T cells. In these cells, HIV expression is silent or reduced, and latently infected cells are thus able to remain undetected by the immune system. Additionally, the pool of latently infected cells can be replenished by clonal expansion in response to antigenic stimulation or homeostatic cues [3–5].

A key mystery in the field is defining the molecular mechanisms that regulate HIV expression and identifying pathways that contribute to maintaining the latent state. The long-lived HIV reservoir is primarily found in resting CD4 T cells, and these cells exhibit low levels of key activating transcription factors (TFs) such as NF-κB, AP-1 and the elongation promoting complex P-TEFb [6–8]. Furthermore, HIV expression in latently infected cells is repressed by the formation of heterochromatin at the integrated provirus, involving a specific set of covalent histone modifications [8–15]. HIV latency is a stable and heritable phenomenon, consistent with the notion of an epigenetic program that regulates HIV latency [16]. In particular, removal of activating marks such as H3K9ac and H3K27ac by class 1 histone deacetylases (HDACs) and the addition of repressive H3K9me3 and H3K27me3 marks by histone methyltransferases have been shown to contribute to HIV latency [17–19]. ATACseq analysis has also shown that latent proviruses exhibit a “closed” chromatin conformation that likely restricts access by key TFs and RNA Polymerase II (RNAPol2) [16].

Latency reversing agents (LRAs) have been developed that targeting various HIV regulating transcriptional pathways, including HDAC inhibitors (HDACis), PKC agonists and non-canonical NF-κB agonists [20]. Some compounds have shown promise for reactivating latent HIV in animal models and in clinical studies [19,21–24]. However, no intervention in persons with HIV (PWH) has yet demonstrated potency for reducing reservoir size. A key issue is that most LRAs are able to reactivate only a fraction of the replication competent reservoir, suggesting that latency is inefficient with single agents [25]. This observation can be explained by a model in which latency is considered as a complex set of states regulated by multiple overlapping mechanisms, and that targeting of multiple pathways concurrently will likely be required to broadly reactivate the reservoir. Thus, fully defining the set of host pathways that impact HIV expression and the key spectrum of limiting pathways in resting CD4 T cells will be crucial for the improvement of latency reversal strategies.

SETD2 is a large (∼290kDa) histone methyltransferase that is crucial for the generation of the H3K36me3 mark in metazoan cells [26]. This enzyme is recruited, in part, by phosphorylated serine-2 on the C-terminal domain of RNA Pol 2 to sites of active transcription and co-transcriptionally deposits H3K36me3 marks in gene bodies [27]. The precise molecular function of this histone mark in gene expression has been the subject of considerable investigation. Current data indicates that H3K36me3 helps to prevent cryptic transcriptional initiation within genes [28], as well as to inhibit repressive histone methyltransferases such as the polycomb repressive complex 2 (PRC2) that generates the transcription-inhibiting H3K27me3 mark [29]. H3K36me3 also regulates DNA repair and transcript splicing [30,31]. However, the role of SETD2 and H3K36me3 in HIV infection is unknown. Significantly, HIV integrase interacts with the PWWP domain containing protein LEDGF, a reader of the H3K36me3 mark, raising the possibility that H3K36me3 could potentially contribute to HIV integration [32]. Indeed, in vitro integration assays demonstrate that H3K36me3 modified histones significantly enhance integration of HIV pre-integration complexes (PICs) into nucleosomal substrates [33,34]. Interestingly, it has also recently been shown that the chromatin insulator CTCF also can mediate HIV integration in microglia and is enriched near regions of high H3K36me3 [35].

Given this connection, we sought to understand whether SETD2 or H3K36me3 contributes to HIV infection by selective inhibition of the enzyme or by CRISPR/Cas9 directed gene knockout. Our results demonstrate that SETD2 or H3K36me3 is not strictly required for HIV infection but plays a key role in regulating post-integration HIV expression and latency. Furthermore, we identify that SETD2s role in HIV expression involves a post-transcriptional mechanism.

## Results

### The SETD2 inhibitor EPZ-719 depletes H3K36me3 in Jurkat cells

To analyze the role of SETD2 activity, we employed the highly potent and selective SETD2 inhibitor EPZ-719 to examine the impact of H3K36me3 loss on HIV infection. We initiated our studies in a human Jurkat CD4 T cell line and observed that exposure of EPZ-719 led to a dose dependent reduction in the level of H3K36me3, with a clear reduction observed by western blot at 100 nM, and a further reduction observable up to 500 nM for a 24h exposure (**Figure 1A**). We also examined the kinetics of H3K36me3 loss in Jurkat cells by taking daily protein samples after 500 nM EPZ-719 exposure. We observed progressive depletion of H3K36me3 over time in the presence of EPZ-719, with the majority of this mark being lost by 24h, and near complete depletion by 72h (**Figure 1B**). These observations confirm that 500 nM exposure of Jurkat cells to EPZ-719 is sufficient to lead to potent reduction of H3K36me3 levels in this cell line.

**Figure 1:**
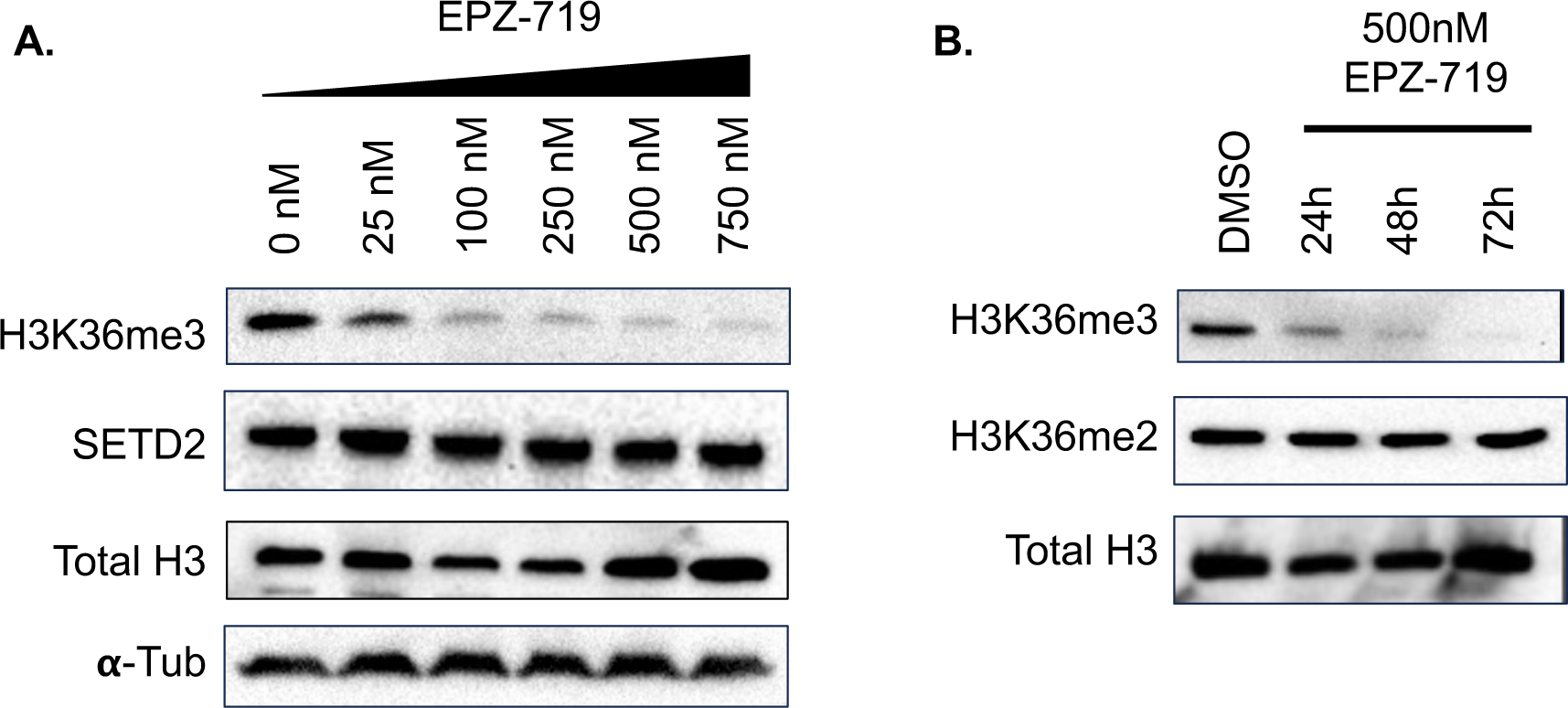
EPZ-719 reduces H3K36me3 levels in human T cell lines. (**A**). Jurkat cells were exposed a range of EPZ-719 concentrations for 24h. Whole cell protein lysates were then extracted and examined by western blot for H3K36me3, SETD2 and total histone 3 (H3) and α-tubulin. (**B**). Jurkat cells were exposed to 500nM EPZ-719 and total cellular protein extracted at times indicated. Extracts were then western blotted for H3K36me3, H3K36me2 and total histone 3 (H3).

### SETD2 activity is not required for HIV infection but regulates HIV expression and latency

To determine the impact of H3K36me3 depletion on HIV infection, we pretreated Jurkat cells with EPZ-719 for three days. We confirmed that these cells were depleted of H3K36me3 by western blot analysis (**Figure 2A**). We then infected the EPZ-719 exposed cells or control vehicle exposed cells with a GFP expressing clone of HIV (HIV-drEGFP). This specific clone contains a short half-life GFP (dreGFP) within the envelope open reading frame, allowing dynamic analysis of LTR-driven HIV expression [17]. When we measured productive HIV infection (%GFP+) by flow cytometry at 3dpi, we observed similar levels of overall infection (29% vs 26%) for EPZ-719 treated cells and cells treated with control vehicle (DMSO) respectively (**Figure 2B, 2C**). These results suggest that SETD2/H3K36me3 is not required for HIV infection. Next, to examine the impact of H3K36me3 on HIV latency, we exposed Jurkat cells to EPZ-719 for 3 days, infected with HIV-dreGFP, then flow sorted GFP+ cells at 3dpi to obtain an enriched actively infected population. We then cultured this infected population in the continuous presence of EPZ-719 or control vehicle for an additional 17 days. During this time, a minor subset of the actively infected cells lost viral gene expression and became GFP-, representing a latently infected population. Interestingly we observed that EPZ-719 exposure significantly impacted the emergence of latently infected cells in the culture. For control treated cells, ∼13% of infected cells cultured in the presence of control vehicle were GFP-by 8 days post infection (dpi). By contrast, EPZ-719 exposed cultures showed a substantially larger fraction of cells that lost GFP expression (∼30%) (**Figure 2D, 2E**), indicating an increased frequency of latently infected cells in the culture. This difference in viral gene expression was maintained at least until 20dpi when the experiment was ended. EPZ-719 did not have any apparent impact on Jurkat cell viability or growth (**Figure S1**). We also examined the impact of EPZ-719 on a spreading infection for a replication competent HIV strain (NL4-3). To study spreading infection, we used a clone of CEM cells (5.25) that contains an integrated LTR-GFP cassette and which becomes GFP+ following productive infection due to *trans*-activation of the reporter cassette by viral expression of Tat [36]. CEM 5.25 cells were pretreated with EPZ-719 for 3 days then infected with NL4-3. After infection with the NL4-3 strain of HIV, we observed rapid viral spread throughout the cell culture indicated by rising GFP+ levels up to 3dpi, followed by a decline as infection saturates. For EPZ-719 exposed cells, spread of the virus was somewhat slower, with reduced levels of GFP+ cells at 2dpi and 3dpi, although ultimately, the virus was still able to spread and overtake the culture (**Figure 2F**). Together, these data are consistent with a model in which SETD2/H3K36me3 is not required for HIV infection but regulates post-integration HIV expression and latency.

**Figure 2:**
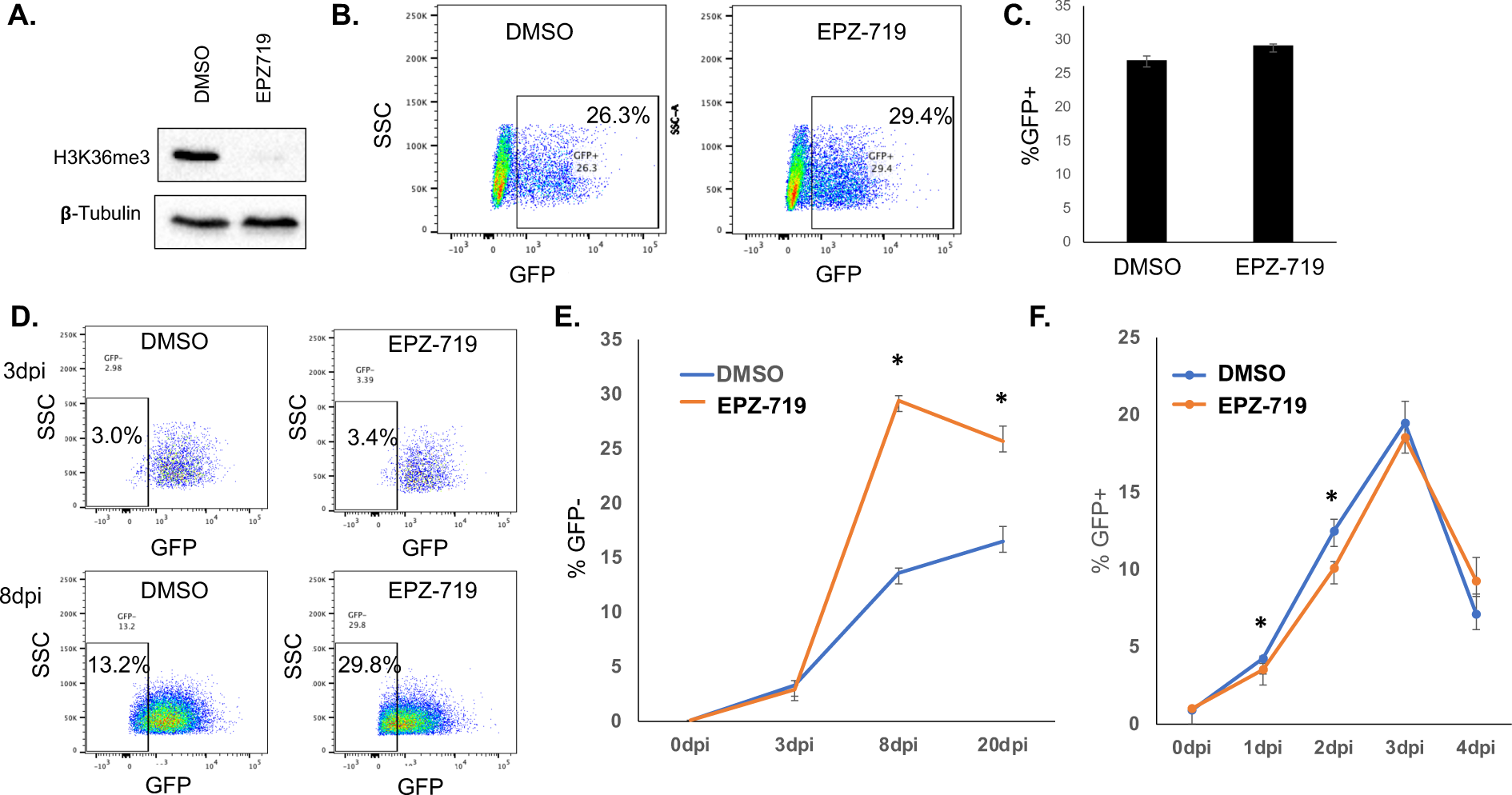
SETD2 activity is not required for HIV infection but regulates HIV expression and latency. (**A**). Jurkat cells were exposed to EPZ-719 for 3 days, then total cell protein lysate was western blotted for H3K36me3 and β-tubulin. (**B**). EPZ-719 treated or control (DMSO) exposed cells (3 days) were infected with HIV-drEGFP, and infection measured at 2dpi by flow cytometry. (**C).** Bar chart of HIV infection for EPZ-719 or DMSO exposed cells. (**D, E**). HIV-dreGFP infected cells were enriched for productively infected cells by flow sporting at 3dpi, then cultured in the presence of EPZ-719 or DMSO for an additional 17 days, and the emergence of a latently infected population (GFP-) was measured by flow cytometry over time. (**F**). CEM5.25 cells were pretreated with EPZ-719 or control (DMSO) for three days, then infected with the replication competent HIV strain NL4-3. Infected cells were then cultured in the presence of EPZ-719 on control for an additional four days. At indicated timepoints, the fraction of infected cells (GFP+) was measured by flow cytometry. Each datapoint represents the average of triplicate samples. Error bars represent the standard deviation of the mean. Asterisk represents a comparison where P<0.05, T Test.

### Inhibition of HIV expression by EPZ-719 is reversible

Since H3K36me3 can directly interact with LEDGF, an HIV integrase interacting factor, we hypothesized that the increased post-integration silencing of HIV in the absence of H3K36me3 might result from a difference in integration sites. If this were the case, then the impact of EPZ-719 on HIV expression from treatment prior to infection should persist even if the drug is removed after viral integration. Similarly, we would predict that HIV expression in infected cells would be resistant to EPZ-719 post integration. We therefore examined whether depletion of H3K36me3 in HIV infected cells induces an ongoing state of transcriptional repression even after withdrawal of the inhibitor. To test this hypothesis, we pretreated Jurkat cells with DMSO control or EPZ-719 for three days to deplete H3K36me3, then infected with HIV-dreGFP, before sorting infected (GFP+) cells. The infected cells were then maintained in EPZ-719 or DMSO for two weeks, to allow a latently infected (GFP-) population to emerge. As expected, EPZ-719 exposed cells exhibited a higher proportion of latently infected (GFP-) cells and reduced levels of H3K36me3 (**Figure 3A, 3B, 3C**). At 2 weeks post infection (wpi), we then divided the culture, and added EPZ-719 to cells that had previously only been exposed to DMSO and removed EPZ-719 from the cells that had been exposed to that, replacing with DMSO. The cells were then maintained in their new condition for 1 week, before re-measuring viral gene expression by flow cytometry (**Figure 3B, 3C**). Notably, when we examined H3K36me3 levels after this 1w exposure, the cells had altered their H3K36me3 levels to match their new condition – newly EPZ-719 exposed cells having lost H3K36me3, and cells that had EPZ-719 removed having restored H3K36me3 levels. In both cases, the infected cells had also altered viral gene expression to match the new exposure – cells that were switched from DMSO to EPZ-719 showed reduced viral expression and elevated latency, while cells that were switched from EPZ-719 to DMSO restored viral gene expression to normal levels. Thus, we conclude that inhibition of HIV expression by EPZ-719 is reversible, along with H3K36me3 levels within the infected cells. This conclusion is inconsistent with inhibition of integration or altered integration site locations contributing to the observed viral suppression in EPZ-719 treated cells. These observations also rule out the possibility that loss of GFP expression in EPZ-719 is due to progressive loss of unintegrated viral DNA over time since this phenomenon would not be reversible.

**Figure 3:**
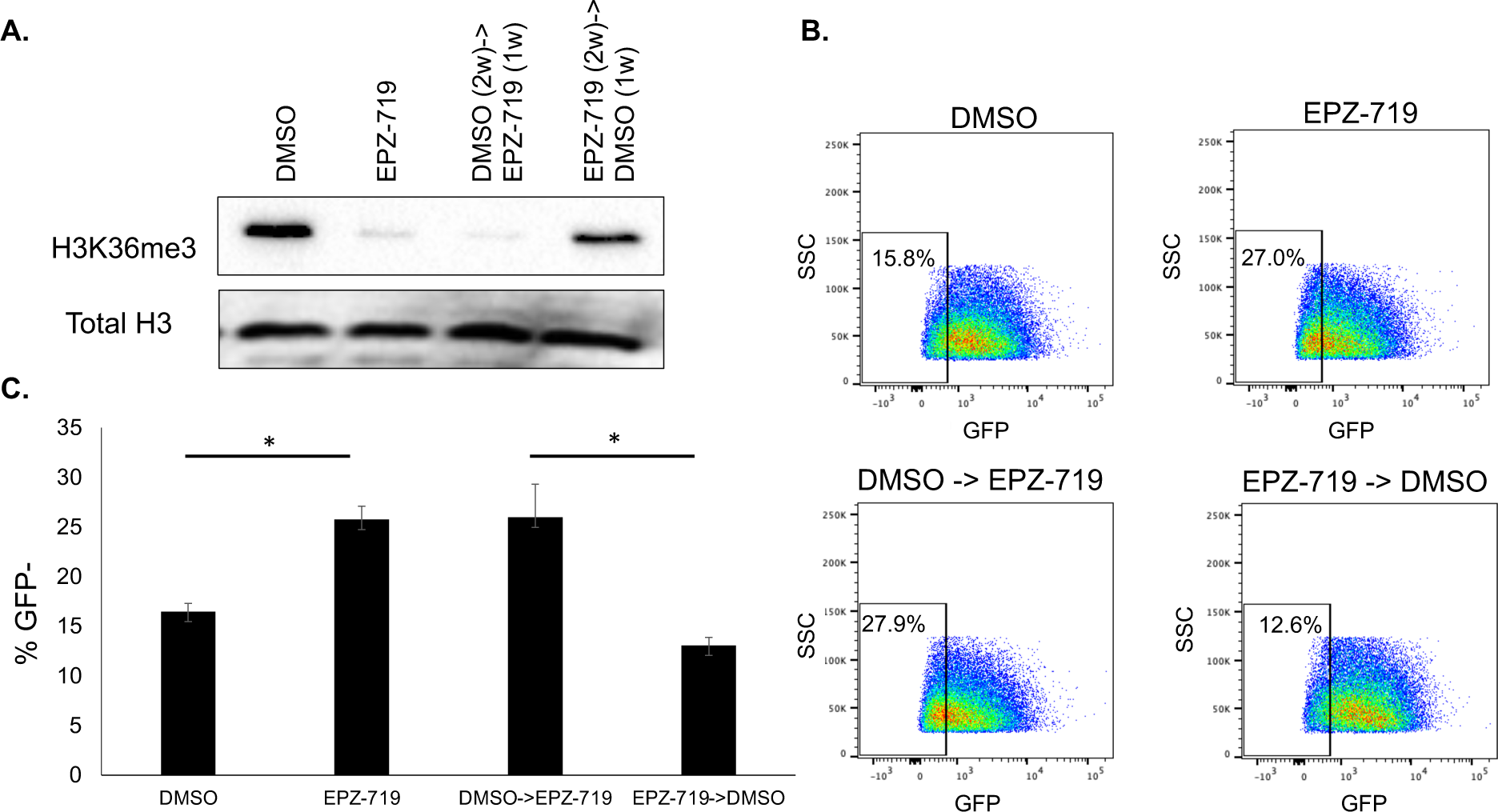
EPZ-719 inhibition of HIV expression is reversible. Jurkat cells were pretreated with EPZ-719 or DMSO for three days, then infected with HIV-dreGFP. At 2dpi, productively infected cells (GFP+) were enriched by flow sorting, then cultured for an additional two weeks in EPZ-719 or DMSO. At 2wpi the culture was divided and exposure conditions were reversed for a subset of the cells before an additional week of culture. At 3wpi, the level of H3K36me3 and total H3 was measured by western blot (**A**), and viral gene expression measured by flow cytometry (**B, C**). Each datapoint represents the average of triplicate samples. Error bars represent the standard deviation of the mean. Asterisk represents a comparison where P<0.05 (T test).

### H3K36me3 loss enhances sensitivity of latent HV to reactivation with an HDAC inhibitor

The presence of latent proviruses is a key barrier to cure for HIV, but reactivation of latent proviruses is typically inefficient. We next decided to examine whether depletion of H3K36me3 from infected cells affects the reactivation of latent HIV using latency reversing agents (LRAs). To test this hypothesis, we exposed Jurkat cells to EPZ-719 for 3 days to deplete H3K36me3, then infected the cells with HIV-dreGFP. Actively infected cells were then enriched by flow sorting for GFP+ cells at 2dpi, then the infected cells were cultured for in continuous DMSO or EPZ-719 for an additional 20 days. After this period of culture, a latent (GFP-) population emerged, and these latently infected cells were further enriched by flow sorting for GFP-cells. Thus, we obtained a polyclonal population of Jurkat cells that were enriched for latently infected cells (**Figure 4A**). Nevertheless, in this population, a subset of cells (28%) still exhibited low level background HIV expression (**Figure 4B**). In the absence of stimulation, the EPZ-719 exposed cells exhibited a modestly lower level of baseline HIV expression (22% GFP+). We then stimulated the latently infected population with two LRAs with distinct mechanisms of action: vorinostat - a class 1 histone deacetylase (HDAC) inhibitor, and prostratin - a protein kinase C agonist. As expected, at 24h post stimulation, both LRAs reactivated a fraction of the latently infected cells, as indicated by an increase in the percent GFP+ cells (**Figure 4B and 4C**), with an average increase of ∼22% GFP+ for vorinostat and an increase of ∼48% for prostratin. Interestingly, while the response to prostratin was similar for the DMSO exposed cells and the EPZ-719 exposed cells, the EPZ-719 exposed cells exhibited a significantly stronger reactivation in response to vorinostat (increase of 44% GFP+ for EPZ-719 exposed cells vs 22% for DMSO exposed cells). To examine further whether this increased response to vorinostat in the absence of H3K36me3, was related to a global elevated responsiveness to HDACi in terms of histone acetylation, we measured total H3K9 and H3K27 acetylation levels for the infected cell population in each of the conditions by western blot (**Figure 4D**). As expected, global H3K9ac and H3K27ac increased significantly in response to vorinostat exposure. Notably, this increase was similar for both EPZ-719 and DMSO exposed cells, suggesting that EPZ-719 does not impact global responsiveness to class 1 HDAC inhibitors, but instead may impact a specific set of genes including HIV.

**Figure 4:**
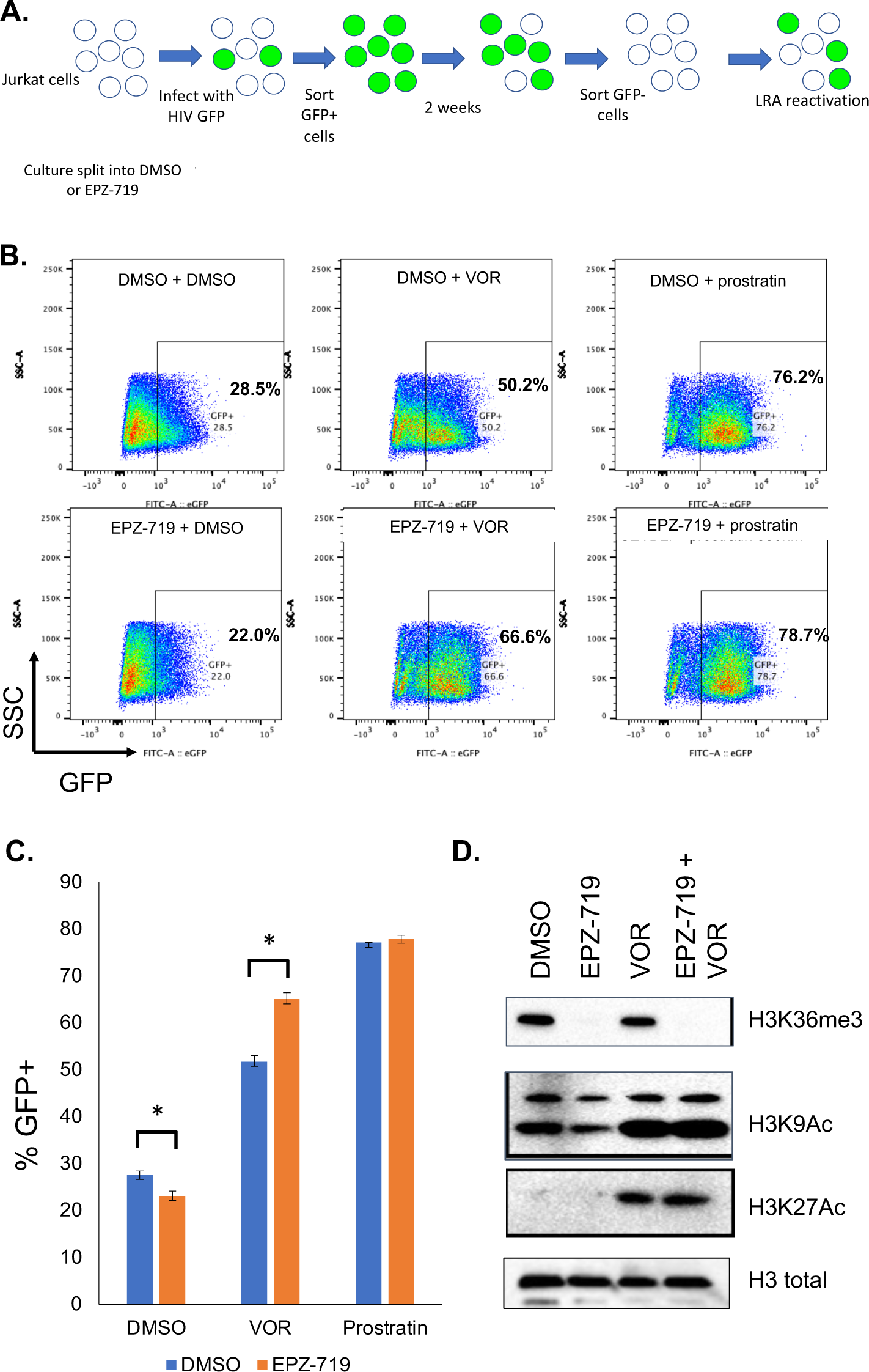
H3K36me3 depletion enhances sensitivity of latent HIV to reactivation with an HDAC inhibitor. (**A**). Schematic overview of experimental design. Jurkat cells were exposed to EPZ-719 (500nM) or DMSO for three days, then infected with HIV-dreGFP. At 48hpi, productively infected cells were isolated by flow sorting, then cultured in EPZ-719 (500nM) or DMSO for two weeks. Latently infected cells (GFP-) were then enriched by flow sorting and stimulated with vorinostat (VOR) (500nM) or prostratin (500nM) in the presence of EPZ-719 (500nM) or DMSO. 24h after stimulation, the reactivation of the latently infected population was measured by flow cytometry. (**B, C**). Flow cytometry and bar chart of HIV-dreGFP reactivation in response to vorinostat (VOR) or prostratin. (**D**). Protein extracts from stimulated and control cells were isolated and histone acetylation (H3K9Ac and H3K27Ac) as well as H3K36me3 was measured for each population by western blot. Each datapoint represents the average of triplicate samples. Error bars represent the standard deviation of the mean. Asterisk represents a comparison where P<0.05 (T test).

### H3K36me3 depletion inhibits HIV expression in primary CD4 T cells

Although Jurkat cells are a valuable model for HIV infection, the natural target cell for HIV is primary CD4 T cells, and several differences exist between primary CD4 T cells and clonal immortalized T cell lines. Thus, we investigated whether our observations regarding the impact of EPZ-719 and H3K36me3 depletion in Jurkat cells could be validated in primary CD4 T cells. We first examined whether exposure of primary CD4 T cells to EPZ-719 led to depletion of H3K36me3. Notably, we observed no significant reduction of H3K36me3 levels in resting CD4 T cells, even after 4 days of exposure (96h) to EPZ-719 (**Figure 5A**). This suggests that, in resting CD4 T cells, SETD2 activity is not required to maintain H3K36me3 over this period of time, likely due to a low rate of removal of this mark and/or low levels of cell division. By contrast, when we added EPZ-719 to recently activated primary CD4 T cells, we observed rapid loss of H3K36me3, with a clear reduction apparent by 24h and near complete loss of H3K36me3 by 48h of exposure (**Figure 5B**). This difference in potency of EPZ-719 between resting and primary CD4 T cells is likely due to more rapid removal of H3K36me3 in activated cells and higher levels of cell division, thereby requiring SETD2 activity to maintain H3K36me3 levels. Thus, we focused on the impact of EPZ-719 on HIV infection in activated primary CD4 T cells. To investigate this, we used a primary CD4 T cell model of HIV infection and latency that we have previously established (**Figure 5C**) [16,17,37]. In this model, primary CD4 T cells are activated for two days then infected with HIV-dreGFP. Actively infected cells (GFP+) are then enriched at 2dpi by flow sorting and cultured for an extended period of time. Infected cells progressively reduce HIV expression as they return to a resting state and latently infected cells (GFP-) emerge over the course of several days. When we combined this model with exposure to EPZ-719 or control vehicle (DMSO) we observed a significant reduction in the percent GFP+ cells for the EPZ-719 exposed cells compared to control cells (22% GFP+ vs 40% GFP+) at 8dpi (**Figure 5D, 5E**). These data indicate that the impact of EPZ-719 and the role of SETD2 in HIV expression is conserved in primary CD4 T cells. In this model, we also examined an inhibitor of the HMT EZH2 (UNC1999) in parallel and in combination with EPZ-719 and found no impact of UNC1999 on HIV expression or on inhibition of HIV by EPZ-719. Thus, inhibition of HIV expression by EPZ-719 was not dependent on EZH2 activity.

**Figure 5:**
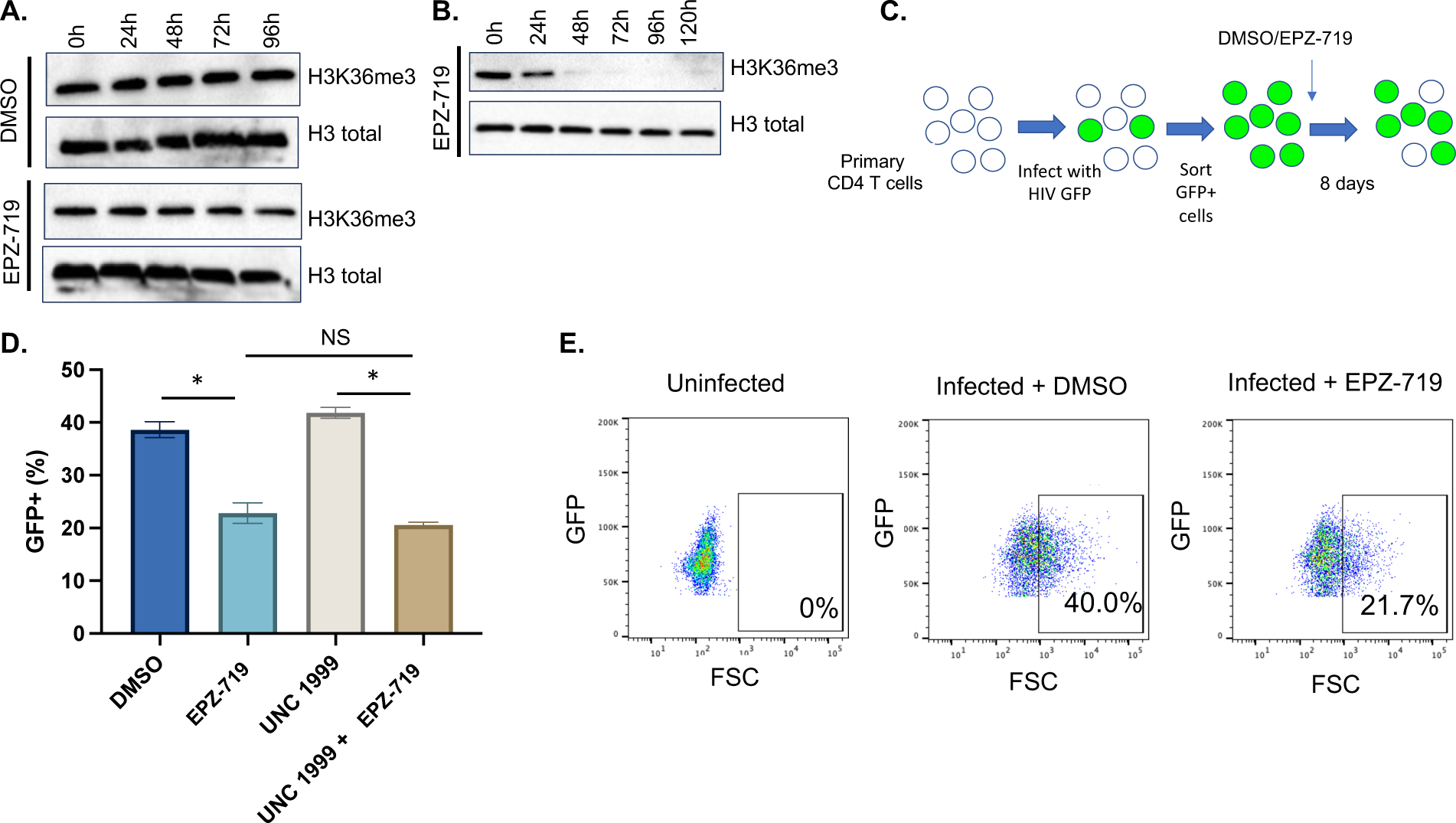
SETD2 activity is required for HIV expression in primary CD4 T cells. 500nM EPZ-719 was added to resting (**A**) or activated (**B**) CD4 T cells for the length of time indicated. Total protein extracts were then harvested and western blotted for H3K36me3 and total histone H3. (**C**) Schematic overview of primary cell model of HIV latency. CD4 T cells were activated for 2d, then infected with HIV-dreGFP in the presence of EPZ-719 (500nM), or DMSO for 48h. GFP+ cells were then flow sorted and cultured for an additional 8 days in the presence of EPZ-719 (500nM), UNC1999 (1μM), EPZ-719+UNC1999 or DMSO. Viral gene expression was then measured by flow cytometry and the percentage of cells with active expression (GFP+) calculated (**D, E**). Each datapoint represents the average of triplicate samples. Error bars represent the standard deviation of the mean. Asterisk represents a comparison where P<0.05 (T Test).

### SETD2 expression is required for HIV expression in primary CD4 T cells

To further confirm the role of SETD2 in HIV expression in primary CD4 cells we performed a CRISPR/Cas9-mediated knockout of SETD2 in HIV infected cells. To achieve this, ribonucleoprotein (RNP) complexes consisting of Cas9 and crRNAs targeting SETD2, the viral transcriptional regulator Tat, or a non-targeting (NT) control were nucleofected into activated HIV-dreGFP infected primary CD4 T cells at 2dpi (**Figure 6A**). Western blot confirmed loss of SETD2 expression in the cells and reduced H3K36me3 levels (**Figure 6B**). When we examined viral gene expression at 8dpi, we observed a significantly reduced level of GFP+ cells in the SETD2 targeted cells compared to cells nucleofected with non-targeting control RNPs. Importantly, targeting of Tat led to a strong reduction in viral gene expression, confirming that this approach can detect regulators of HIV expression (**Figure 6C, 6D**). Overall, these data provide further confirmation that SETD2 is an important regulator of HIV expression in primary CD4 T cells and that SETD2 activity is required to maintain active HIV gene expression in CD4 T cells.

**Figure 6:**
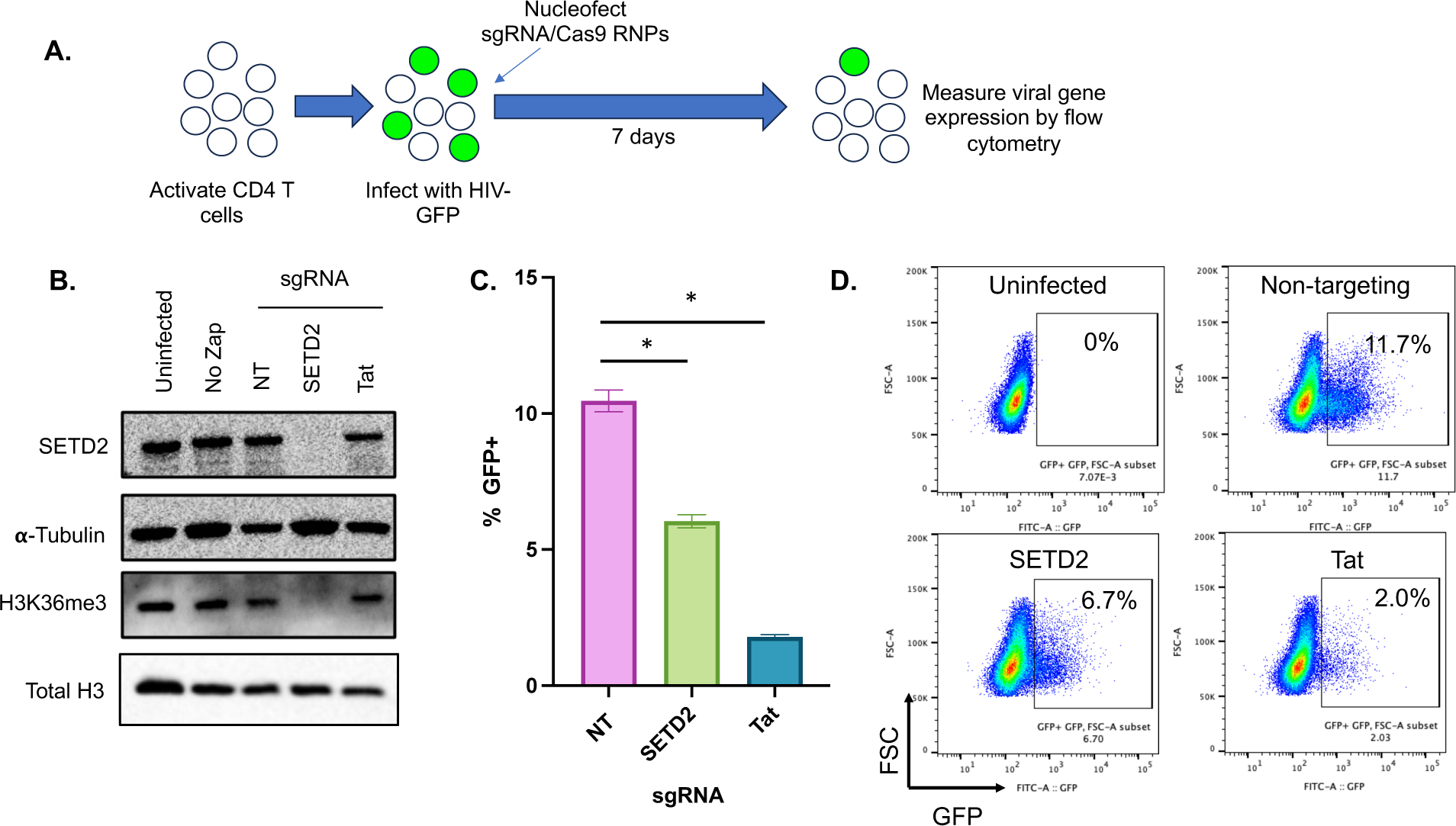
SETD2 knockout in primary CD4 T cells reduces HIV expression. (**A**). Schematic overview of SETD2 knockout in primary CD4 T cells. CD4 T cells were activated for 2 days, then infected with HIV-dreGFP. At 2dpi, cells were nucleofected with ribonucleoparticles (RNPs) targeting SETD2, Tat, or non-targeting (NT) control. At 8d post targeting total cell protein lysate was extracted and western blotted for SETD2, H3K36me3, total histone H3 and α-tubulin (**B**). Uninfected cells and infected but not nucleofected cells (“No Zap”) are also shown. Viral gene expression was measured by flow cytometry (**C,D**). Each datapoint represents the average of triplicate samples. Error bars represent the standard deviation of the mean. Asterisk represents a comparison where P<0.05 (T test).

### EPZ-719 affects cellular gene expression but does not alter HIV RNA levels

To further understand the role of SETD2 in regulation of HIV expression, we performed RNAseq on HIV infected Jurkat cells that had been cultured in the presence of EPZ-719 or control vehicle (DMSO). Interestingly, we observed many differentially expressed genes (DEGs) between the two conditions, 4833 genes total (p_adj_<0.05), with 2310 genes upregulated and 2523 downregulated (**Figure 7A, Table S1**). However, most of these changes were modest in magnitude with only 34 upregulated DEGs and 38 downregulated DEGs changing by greater than log_2_FC>1. Nevertheless, we observed some intriguing patterns in the transcriptomic changes. When we examined upregulated genes for enrichment with particular chromatin features using Enrichr [38], we observed a strong enrichment for genes that are associated with E2F binding (**Figure 7B, upper panel, Table S2**). By contrast, downregulated genes were highly associated with Myc binding peaks (**Figure 7B lower panel, Table S3**). Notably Myc expression itself was reduced by EPZ-719 exposure, consistent with an overall reduction in Myc-dependent gene expression. We also performed gene ontology (GO) analysis [39] of the top 500 upregulated or downregulated genes and observed that upregulated DEGs were highly enriched for structural constituents of chromatin (GO:0030527) (pval_adj_= 7.96×10-14) while downregulated genes were highly enriched in RNApol2 specific DNA binding transcription factor activity (GO:0003700, pval_adj_= 3.55E-20) (**Tables S4-S7**). These observations are consistent with EPZ-719 having a broad effect on cellular chromatin and RNApol2 driven transcription.

**Figure 7:**
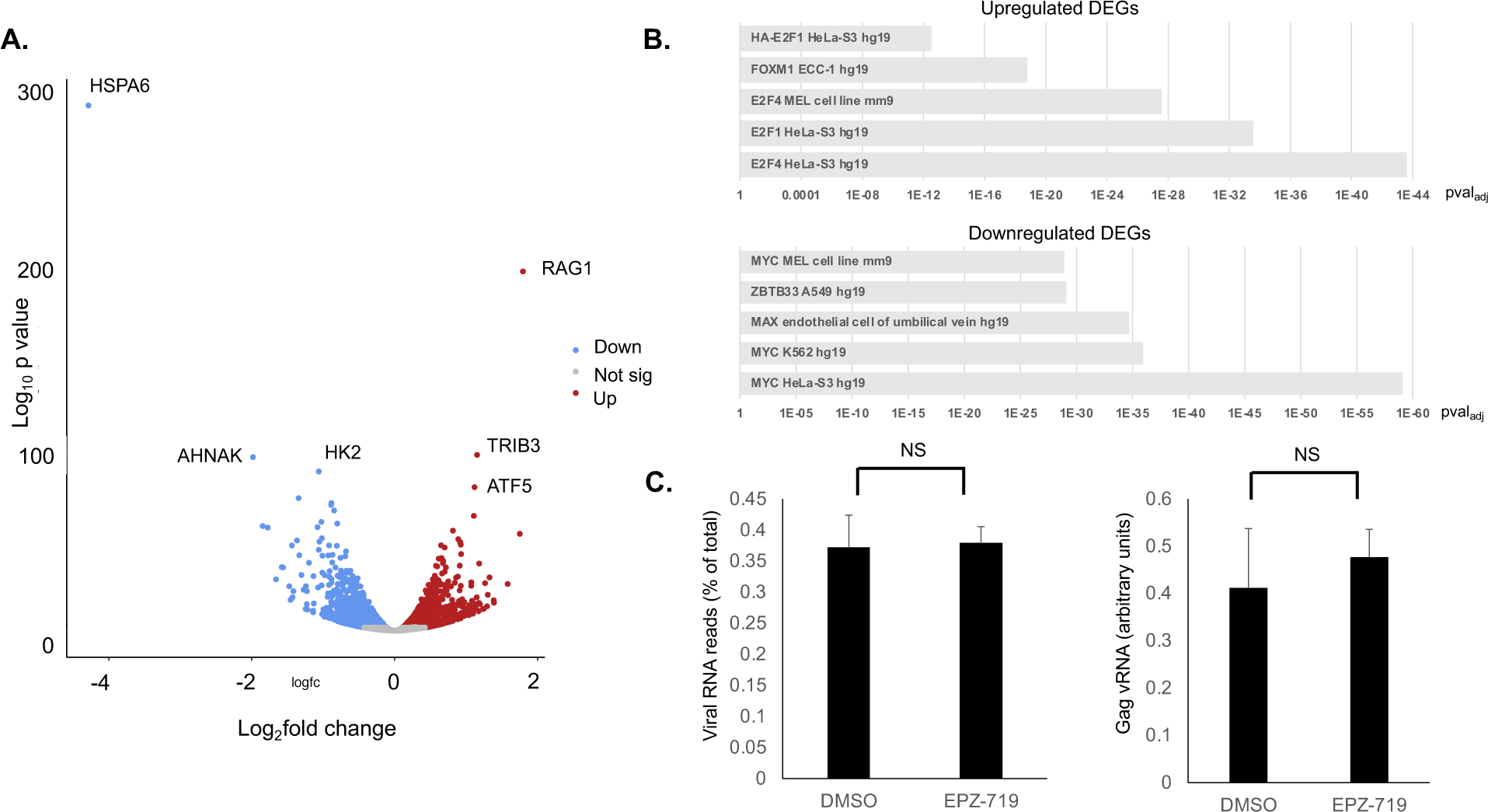
EPZ-719 affects cellular gene expression but not HIV RNA levels. HIV-dreGFP infected Jurkat cells that had been exposed to EPZ-719 (500nM) or DMSO for three days before infection and 8 days after infection, were profiled by bulk RNAseq. **A.** Volcano plot of differentially expressed genes (DEGs), with significantly downregulated (pval_adj_<0.05) genes in blue and upregulated genes shown in red. The three most significantly upregulated and downregulated genes are labeled. Not sig= not significantly different (grey). **B.** The upregulated and downregulated genes were analyzed using Enrichr to find enrichment of known protein/DNA binding sites from the ENCODE databse with the gene set. **C.** The abundance of unique viral reads as a percentage of total unique reads in the dataset was calculated (left panel), and the abundance of Gag viral RNA (Gag-vRNA) was also examined by quantitative PCR (right panel, arbitrary units). The analysis was performed on biological triplicate samples. Error bars represent the standard deviation of mean. NS= not significant, T test (P<0.05).

We also examined expression of several known regulators of HIV transcription in the RNAseq dataset. We previously identified the chromatin insulator CTCF as inhibiting HIV expression in infected cells [16], and CTCF expression was upregulated by EPZ-719 exposure. Expression of the catalytic subunit of the Polycomb Repressive Complex 2 (PRC2) EZH2 was also increased in EPZ-719 exposed cells. Interestingly, we observed some evidence of innate immune activation in EPZ-719 exposed cells – expression of the interferon regulated genes IRF1, IRF4, MX2, IFIT3, and IFIT2 were all upregulated. Additionally, cytokine signaling mediators STAT4 and STAT2 were both upregulated in response to EPZ-719. We also examined the expression of HIV restriction factors in the dataset and observed mixed outcomes. APOBEC3F, APOBEC3D, as well as APOBEC2 were all upregulated in the presence of EPZ-719. By contrast, BST2, SERINC1, SERINC2, and SERINC5 were downreglated. Overall, these data indicate that EPZ-719 affects a broad number of cellular genes, including several that could have a direct or indirect impact on HIV expression.

To investigate whether EPZ-719 affects the HIV RNA expression level, we also compared the overall fraction of HIV mapping reads in the EPZ-719 exposed cells to the control cells. Interestingly, the abundance of HIV reads was similar between the two conditions indicating no impact of EPZ-719 on overall HIV RNA levels (**Figure 7C, left panel**). To confirm this observation, we carried out quantitative PCR analysis of Gag RNA levels in HIV-dreGFP infected cells that had been cultured in the presence of EPZ-719 or DMSO for 8 days (**Figure 7C right panel**). Similar to the RNAseq results, we found no significant difference in the levels of Gag RNA with EPZ-719 exposure, although we did observe a trend towards increased Gag RNA in EPZ-719 exposed cells relative to control exposed cells. This observation suggests that EPZ-719 does not reduce the overall level of HIV transcription and RNA abundance, but instead may be impacting a post-transcriptional step in HIV expression.

### Impact of EPZ-719 on HIV integration sites

To further examine whether SETD2 activity or H3K36me3 levels in target cells impacted HIV integration, we extracted cellular genomic DNA from Jurkat cells that been exposed to EPZ-719 or control (DMSO) for 3 days then infected with HIV-dreGFP. Integration sites from each condition were then determined by linear amplification from the HIV LTR followed by nested PCR and long read sequencing of the amplified product [40]. Using this approach, we recovered a similar number of integration sites from cells in either the DMSO or EPZ-719 condition (781 and 777 respectively). When we examined the chromosomal distribution of the integration sites, we observed, consistent with previous reports, that distribution across the chromosomes was non-random, with certain chromosomes such as Ch19 having elevated enriched abundance of integration sites (**Figure 8A)** [41]. When we compared integration sites between cells infected in the presence of EPZ-719 to cells infected in the presence of DMSO, we observed an overall similar distribution across the chromosomes, with only minor differences apparent (**Figure 8A**). We then examined the association of integration sites with gene regions within our dataset. As expected, the majority of HIV integration sites (79%) were associated with genes rather than intergenic regions (21%) for cells infected in the presence of DMSO (**Figure 8B**). This observation was also true for EPZ-719 exposed cells, with 74% of integration sites associated with genes and 26% within intergenic regions. However, we observed a statistically significant (p<0.05, T test) increase in integration in non-genic regions in the presence of EPZ-719. We conclude from this that the association of HIV integration with genes does not require SETD2 activity or H3K36 methylation, but that SETD2 activity does modestly affect the distribution of integration sites.

**Figure 8:**
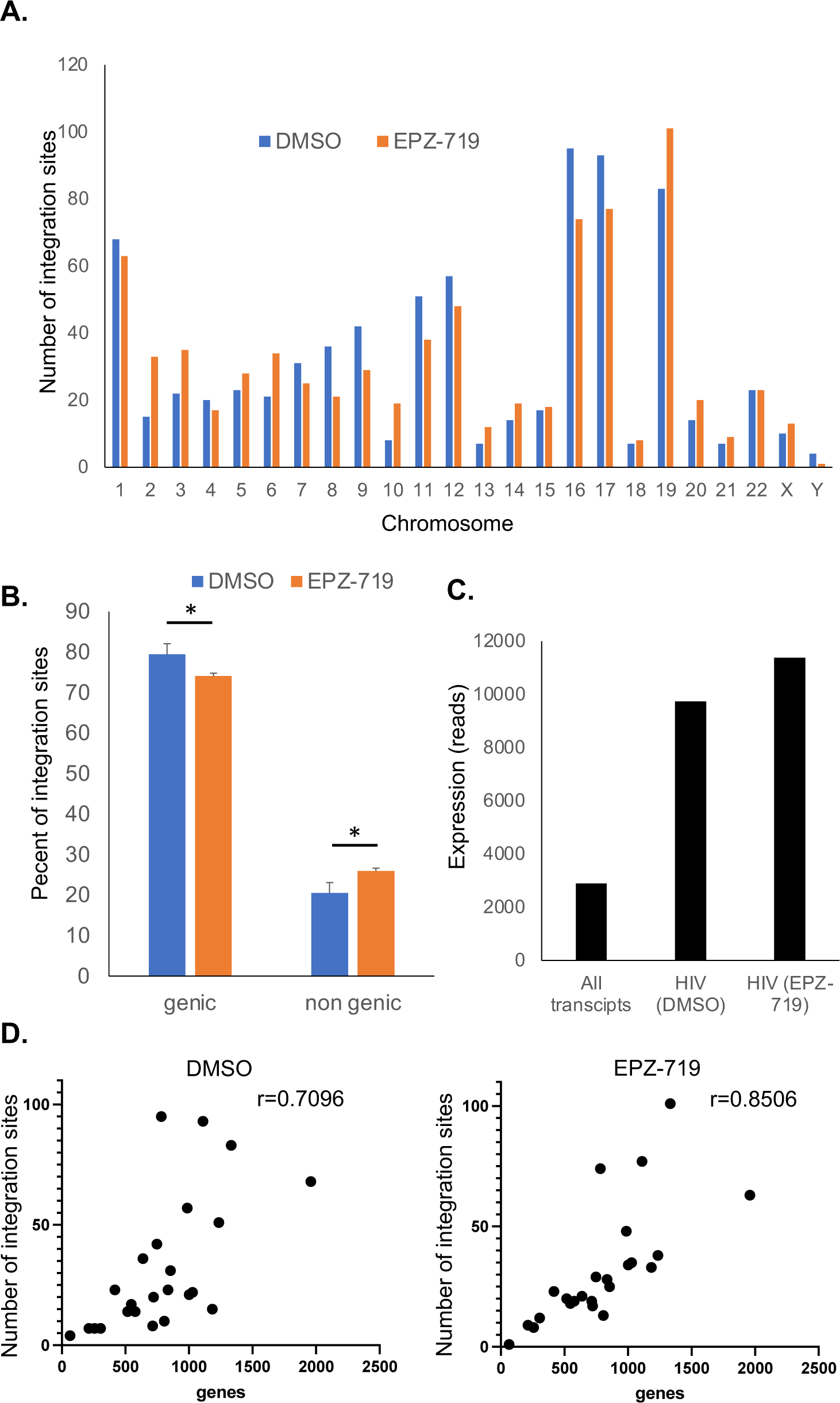
Analysis of HIV integration sites in the presence of EPZ-719. Genomic DNA derived from Jurkat cells infected with HIV-dreGFP in the presence of EPZ-719 (500nM) or control (DMSO) as used to identify 777 and 781 integration sites respectively. For these integration sites, we compared their chromosomal distribution (**A**), their association with genes (**B**), the normalized expression level of cellular genes associated with HIV integration sites that were infected in the presence of EPZ-719 or DMSO control (**C**), and the Spearman correlation between the number of genes and the number of integration sites across chromosomes (**D**).

Since H3K36me3 is deposited by SETD2 during transcription and is associated with actively expressed genes, we also examined the expression level for genes that were associated with HIV integration sites in the presence of EPZ-719 or DMSO control. For both conditions, HIV integration was enriched in genes that were highly expressed in infected cells – the average normalized transcript read count for all genes was 2890, while, for genes associated with HIV integration sites in EPZ-719 or DMSO treated cells, the average expression level was 11337 and 9739 respectively (**Figure 8C**). Thus, EPZ-719 exposure and H3K36me3 depletion did not affect the association of HIV integration with highly expressed genes. Additionally, we examined the correlation between the number of integration sites on each chromosome and the number of genes on that chromosome. We observed a strong positive correlation between the number of genes on a chromosome and the number of integration sites. This observation was true for both control cells and EPZ-719 exposed cells (r= 0.7096 and r= 0.8506 respectively) (**Figure 8D**). Overall, these results indicate that EPZ-719 exposure does not have a major effect on viral integration sites.

### SETD2/H3K36me3 affects HIV RNA splicing

We next considered post-transcriptional mechanisms by which EPZ-719 might affect HIV expression. H3K36me3 has been shown to regulate m6A modification of RNA by METTL3 [42], and m6A RNA methylation significantly impacts HIV expression and replication [43–46]. We therefore examined total cellular m6A levels in cellular RNA after 13d of exposure of HIV-dreGFP infected cells to 500nM EPZ-719 (**Figure S2A**). After this period of culture, EPZ-719 exposed cells exhibited reduced HIV expression as expected. However, despite a clear difference in the expression of HIV and the abundance of latently infected (GFP-) cells in the culture (**Figure S2B**), we observed no change in total m6A abundance compared to DMSO control exposed cells (**Figure S2C**). To examine m6A RNA modification within HIV transcripts, we also performed immunoprecipitation of m6A RNA from infected cells, followed by quantitative PCR for a region of HIV within the Env/Rev coding sequence that has been shown to be abundantly modified with m6A [44,45]. As expected, m6A-specific pull down significantly enriched this RNA sequence over control IgG, but we observed no difference in abundance between EPZ-719 exposed cells and control (DMSO) exposed cells (**Figure S2D**). Thus, we conclude that EPZ-719 likely does not impact HIV expression through affecting m6A modification of viral or cellular RNAs.

We next considered whether SETD2 might be impacting HIV transcript splicing. The full-length HIV genomic transcript can be extensively spliced to generate up to 50 different RNA forms, using four splice donor sites (D1, D2, D3, D4) and ten different acceptor sites (A1, A2, A3, A4a, A4b, A4c, A4d, A5, A5b, A7) (**Figure 9A**). To investigate whether EPZ-719 affects viral splicing, we used a recently developed assay to quantify individual HIV RNA species in HIV-dreGFP infected cells after 13d of exposure to EPZ-719 at 500nM [47]. This assay involves first performing a reverse transcription reaction with a random unique molecular identifier that also serves as a primer to capture all HIV RNA species - unspliced, 4kb, and 1.8kb. During reverse transcription this random unique molecular identifier is added to the cDNA, allowing accurate quantification of each molecule in the original pool after subsequent PCR amplification and Illumina sequencing. After performing this analysis, we observed that in the EPZ-719 treated cells, there was a significant overall reduction in the fraction of spliced transcripts compared to the control cells (59% vs 69%) (**Figure 9B**). Additionally, for the EPZ-719 treated cells, there was a small reduction in the percentage of 1.8kb transcripts as a fraction of the sliced transcripts (**Figure 9C**). Furthermore, when we examined splice donor and acceptor usage within the spliced transcripts, we noticed an increase in usage of the A1 initial acceptor and a reduction in the A5 initial acceptor usage (**Figure 9D**). However, when we compared the final acceptor usage, there was no significant difference between the two conditions. These observations indicate that loss of H3K36me3 can alter viral splicing in infected cells (**Figure 9E**) and leads to a reduction in spliced viral RNAs. This effect on viral splicing may contribute to the overall suppressive effect of EPZ-719 on viral expression.

**Figure 9:**
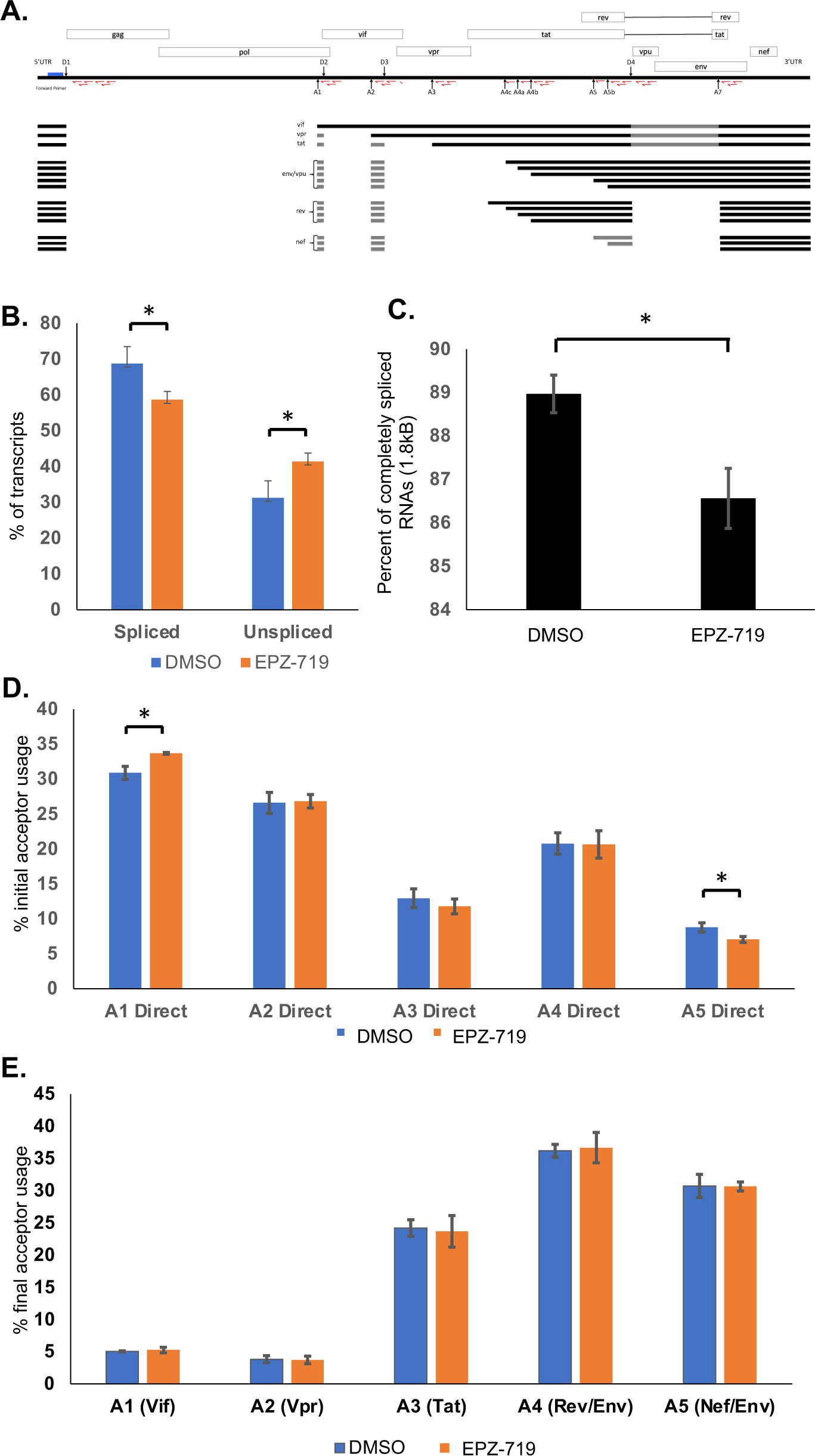
EPZ-719 affects HIV RNA splicing. HIV RNA splice variants were quantified by a method in which primers including a random barcode sequence of nucleotides were used for reverse transcription, followed by PCR for the major 4kb and 1.8kb species of viral RNA, and Illumina sequencing. (**A**). The location of the primer binding sites and the HIV slice donor (D1-D4) and acceptor (A1-A7) sites are shown on a map of the HIV genome. (**B**). The fraction of vRNAs that are spliced is shown. (**C**). The percentage of completely spliced vRNAs (1.8kb) is shown. (**D**). The fraction of spliced transcripts using different initial acceptors is shown. (**E**). The fraction of spliced transcripts using different final acceptors is shown. Each bar represents the average of three independent biological replicates. Error bars represent the standard deviation of the mean. Asterisk indicates significant difference (P<0.05, Students T test.

## Discussion

Defining the set of host factors that regulates HIV expression will be essential for developing approaches to broadly reactivate the latent HIV reservoir. Studies have shown that HIV expression is regulated by an array of host cell transcriptional complexes and chromatin modifying enzymes [48,49]. Curiously, latent HIV proviruses retain features of transcriptionally repressed heterochromatic regions despite HIV integration occurring primarily in actively transcribed host genes. Since many of these chromatin features can be vertically transmitted during cell division, latency likely represents a stable and heritable epigenetic state. Thus, in addition to being important for direct repression of HIV expression within a cell, latency-associated chromatin likely plays a key role in reservoir persistence by transmitting the latent state during clonal expansion of infected cells in people with HIV [10,14,50,51]. Histone-modifying enzymes have been examined for specific roles in HIV latency and reactivation and various complexes have been found to play positive or negative roles in HIV expression. In particular, the role of histone deacetylases (HDACs) in the establishment and maintenance of latency is well known, and HDAC inhibitors potently reverse and prevent HIV latency, both in cell-based model systems and in PWH [17,19,52–55]. The histone methyltransferase EZH2, the catalytic subunit of the polycomb repressive complex 2 (PRC2), may also play a role in HIV silencing [11,56–58]. More recently, the HMTs SMYD2 and SMYD5 have been implicated in HIV expression and latency. SMYD5 has been shown to be required for high HIV expression and promotes HIV transcription by binding to the TAR RNA and mediating methylation of Tat [59,60].

Notably, the role of the SETD2 in HIV infection has not been previously examined. SETD2 is a large (288kDa) protein that is abundantly expressed in mammalian cells and is solely responsible for generating H3K36me3 [27]. The functional role of H3K36me3 has previously been examined and its overall role in transcription and gene expression appears to be complex. H3K36me3 is highly enriched in the gene bodies of actively transcribed genes and has been shown to repress cryptic transcriptional initiation sites within genes through the recruitment of HDACs and other repressive modifications, including DNA methylation [28,61,62]. Other data suggests that H3K36me3 limits the spread of the repressive H3K27me3 mark within actively expressed genes [63] in addition to modulating post-transcriptional events that include RNA processing and m6A methylation of RNA [42]. Notably, SETD2 has been shown to play a role in viral replication. The SARS-Cov2 protein NSP9 inhibits the interferon response pathway through targeting SETD2 [64], and SETD2 has also been implicated in transcriptional regulation of human papilloma virus (HPV) [65,66]. As such, in this manuscript we have investigated the role of SETD2 in HIV gene expression in latency.

We found in this work that SETD2 expression and its activity is important for promoting high HIV expression in infected cells and for limiting the emergence of a latently infected cell population. By using a selective inhibitor of SETD2 or by CRISPR-mediated knockout of SETD2 we show that loss of SETD2 and H3K36me3 was associated with significantly reduced HIV expression and an increased fraction of cells with a latent viral phenotype. This finding was confirmed in both Jurkat cell and primary cell models of HIV infection. Notably H3K36me3 depletion did not have an impact on the overall number of infected cells after adding HIV virus particles to target cells, indicating that SETD2 and H3K36me3 are not required for HIV infection and integration but affect a post-integration aspect of viral gene expression. We had initially hypothesized that H3K36me3 might contribute to HIV integration, since the integrase associated factor LEDGF binds to H3K36me3, and HIV integration sites are enriched in H3K36me3 dense regions. However, we observed that HIV integration sites were largely similar in the presence or absence of H3K36me3, with a similar chromosomal distribution and preference for highly transcribed genes. Nevertheless, we did observe a modest shift in the integration sites towards non genes in the absence of H3K36me3. Given that the H3K36me2 mark generated by NSD enzymes is also predicted to recruit LEDGF, we speculate that H3K36me2 plays a compensatory role in the absence of SETD2/H3K36me3.

Interestingly, depletion of H3K36me3 also increased the sensitivity of latent proviruses to reactivation with the histone deacetylase inhibitor vorinostat. The mechanism behind this phenomenon is unclear, but previous has work has shown that inhibiting the HMT EZH2 can also enhance the responsiveness of latent proviruses to vorinostat [57]. H3K36me3 represses transcriptional initiation within gene bodies, but it is also possible that, in some infected cells, H3K36me3 is also found at the viral LTR where it could contribute to suppression of HIV transcription. Although we found that SETD2 inhibition alone did not impact HIV transcription, removal of LTR-associated H3K36me3 could nevertheless render a subpopulation of viral promoters in a poised state that responds to HDAC inhibition, in addition to a post-transcriptional effect. Depleting H3K36me3 with EPZ-719 may also help to improve the responsiveness of the clinical reservoir to vorinostat. However, an obstacle to this approach is the apparently limited efficacy of EPZ-719 in resting CD4 T cells. This low activity likely results from slow turnover of H3K36me3 in resting CD4 T cells, and it is possible that enhancing demethylation of this residue could achieve a similar effect to inhibiting SETD2.

Based on our findings we propose that the effect of SETD2 on HIV expression is primarily at the post-transcriptional level. Despite robust changes to HIV expression at the protein level in EPZ-719 exposed or SETD2-depleted cells, we observed no change to the overall level of HIV RNA in either RNAseq data or in qPCR for viral RNA. SETD2 has previously been shown to affect several aspects of post-transcriptional RNA processing. In particular, SETD2 was shown to be required for m6A methylation of cellular RNA, and this RNA modification is known to strongly impact gene expression. Other labs have also previously shown that the HIV RNA is extensively methylated with m6A [43–46]. Nevertheless, we did not observe any effect of SETD2 on total or HIV m6A RNA methylation in our study. It remains possible that SETD2 affects a different less well characterized form of RNA modification that is required for efficient HIV expression.

We also examined HIV RNA transcript splicing and found that SETD2 inhibition affected the fraction of spliced transcripts. Specifically, overall HIV splicing was significantly reduced in the presence of EPZ-719. Furthermore, within spliced transcripts some modest differences in donor/acceptor usage were also observed. These data suggest that the presence of SETD2 and H3K36me3 on HIV provirus-associated histones during transcription affects post transcriptional maturation of viral RNA. It is possible that the altered transcript splicing pattern for HIV in the absence of SETD2 activity contributes to the reduced overall level of viral gene expression. However, it remains possible that other post transcriptional mechanism also contribute to SETD2’s role in HIV expression.

While we propose a lack of H3K36me3 at the HIV locus results in defective splicing of HIV that leads to its overall reduced expression, it is also possible that SETD2 exerts an indirect effect on HIV expression. Indeed, global transcriptomic profiling of EPZ-719 exposed cells indicated altered expression of many cellular genes. Interestingly, increased expression of several interferon sensitive genes (ISGs) was observed when SETD2 activity was inhibited, suggesting induction of an overall antiviral state that could contribute to inhibiting HIV expression. Notably, previous work has linked SETD2 to the innate immune system. Specifically, SETD2 has been shown to amplify IFN signaling by methylating STAT1 [67]. It is important to note that SETD2 has been suggested to methylate other cellular targets other than H3K36, which could conceivably be important for regulation of HIV. For example, in addition to STAT1, SETD2 has been suggested to trimemethylate actin on K68 [68].

Finally, a key question raised by our findings is whether loss of SETD2 activity or H3K36me3 depletion is associated with the formation of maintenance of the clinical reservoir. Variation in SETD2 expression or activity and/or abundance of H3K36me3 could impact whether a given infected cell adopts a latent or active infection phenotype. Additional studies will be needed to demonstrate the role of SETD2 in regulation of the clinical reservoir and its potential as a target to enhance latency reversal in vivo.

## Methods and Materials

### Cell lines

HEK293T cells were maintained in DMEM media with 10% fetal bovine serum (FBS) and penicillin/streptomycin. Jurkat cells, CEM5.25 cells and primary CD4 T cells were maintained in RPMI media with 10% FBS, penicillin/streptomycin, and supplemented with glutamine, HEPES and sodium pyruvate. 2D10 cells were a gift from Dr Jonathan Karn. CEM5.25 cells were provided by Dr Dan Littman. 293T cells were obtained from ATCC.

### Virus infections

HIV particles were produced by transfection of HEK293T cells with viral plasmid DNA using Mirus LT1 reagent (Mirus Bio LLC, Madison WI). At 24h, supernatant was replaced with fresh RPMI to collect virus. At 48h, viral supernatant was collected. Cellular debris was removed first by low-speed centrifugation (300g, 5min), then filtration through at 0.45μm filter. The clarified supernatant was then frozen in aliquots and titered by infecting Jurkat cells with a dilution series of virus supernatant followed by flow cytometry for GFP+ cells at 48h. For the HIV-dreGFP virus (NL4-3-D6-dreGFP/Thy1.2), viral backbone plasmid was co-transfected with two packaging plasmids – PAX2 (encoding HIV GagPol) and MD2-VSVG (encoding the VSV-G glycoprotein). NL4-3-D6-dreGFP/Thy1.2 was modified from a parental plasmid provided by Dr R Siliciano. The replication competent NL4-3 strain clone was obtained from Aidsreagent.org.

### Flow cytometry and cell sorting

Samples of infected cells were analyzed using a Fortessa flow cytometer (Becton Dickson). Viable cells were gated using a live/dead stain – Zombie Violet (Biolegend) to stain cells prior to analysis. Doublet cells were excluded using forward scatter and side scatter. For flow sorting to enrich cells based on GFP expression, a FACSAria machine (Becton Dickson) in BSL2+ containment facility was used.

### Primary cell model of HIV latency

Human whole blood was obtained from StemCell Inc (Vancouver), and total CD4 T cells isolated using a negative selection CD4 T cell enrichment kit (StemCell). Aliquots of CD4 T cells were frozen and stored in 90% FBS 10% dimethylsulfoxide (DMSO). Aliquots of frozen cells were then thawed and activated with anti-CD3/CD28 beads for 48h (Life Technologies). Activated CD4 T cells were then spinocculated with HIV-dreGFP virus supernatant for 2h at 600g, before plating in RPMI with IL-2 (100U/mL) and IL-7 (5ng/mL). Infection was measured at 48h post infection by flow cytometry. Cell media was replaced every 2-3 days, and the cells were maintained at a density of between 1-2 million per mL. In some experiments, GFP+ cells were sorted at 48 hours post infection to enrich for actively infected cells using a FACSAria (Becton Dickson).

### CRISPR-Cas9 gene knockouts

CRISPR RNA targeting sequences (crRNA) against SETD2 were designed using CRISPick [69], while Tat-targeting and non-targeting control sequences were derived from previous literature [70,71]. CRISPR-Cas9 ribonucleoprotein complexes (RNPs) were prepared by mixing crRNA targeting sequence and tracrRNA (IDT) at 1:1 to a final concentration of 100μM and annealed in duplex buffer (IDT) by heating to 95°C and slow cooling to room temperature in a PCR thermocycler. RNP complexes were generated immediately before nucleofection by mixing 0.8μL of Cas9 enzyme (62μM concentration, 49.6pMol total, ALT-R by IDT), 0.8μL of poly-glutamic acid (15kDa average polymer size, 100mg/mL concentration, Alamanda laboratories), and 1μL of annealed duplex crRNA:tracrRNA (100pMol for a final molar ratio of approximately 2:1 RNA:Cas9 enzyme). For SETD2 targeting, three crRNAs were multiplexed for more efficient target knockout. Primary CD4 T cells were isolated, activated, and infected as above. Six days after activation (4 days post infection), 3×10^6^ cells per condition were pelleted at 90g for 10 minutes, washed in PBS, and then resuspended in primary cell nucleofection buffer P3 (Lonza) at 1×10^8^ cells per mL (2×10^6^ cells per 20μL reaction volume). Cell suspensions were then added to RNP complexes, and nucleofection was performed using the CM137 protocol (Lonza 4D nucleofector). Nucleofected cells were then immediately resuspended in pre-warmed RPMI and incubated at 37° for 20 min prior to resuspension at 2×10^6^ cells per mL in complete RPMI with IL-2 and IL-7. Specific crRNA sequences used were:

SETD2

AltR1/rGrG rArGrU rCrGrA rGrUrC rUrArC rCrUrG rArArG rGrUrU rUrUrA rGrArG rCrUrA rUrGrC rU/AltR2/

/AltR1/rGrC rUrCrA rArGrG rUrGrA rArArU rArGrC rArUrG rGrUrU rUrUrA rGrArG rCrUrA rUrGrC rU/AltR2/

/AltR1/rArU rGrArA rCrUrG rGrGrA rUrUrC rCrGrA rCrGrA rGrUrU rUrUrA rGrArG rCrUrA rUrGrC rU/AltR2/

Non-Targeting(NT)

/AlTR1/rArCrGrGrArGrGrCrUrArArGrCrGrUrCrGrCrArArGrUrUrUrUrArGrArGrCrUrArUrGrCrU/A lTR2/

Tat

/AlTR1/rCrCrUrUrArGrGrCrArUrCrUrCrCrUrArUrGrGrCrGrUrUrUrUrArGrArGrCrUrArUrGrCrU/A lTR2/

### Cellular fractionation and western blotting

To produce protein lysates, cells were pelleted by centrifugation (300g 5min), then washed in phosphate buffered saline (PBS), before being lysed in RIPA buffer (ThermoFisher) supplemented with protease inhibitors (Roche). Cell protein lysates were quantified by Bradford assay (ThermoFisher), then separated by Tris-Glycine SDS-PAGE, and transferred to polyvinylidene difluoride (PVDF) or nitrocellulose membranes. Blots were blocked in 5% milk in Tris-Buffered Saline (TBS), then incubated overnight with primary antibodies diluted in TBS containing 5% bovine serum albumin (BSA) or 5% milk and washed with 0.1% Tween20 TBS (TBST). Membranes were then stained with secondary antibodies (1:10,000) conjugated to horseradish peroxidase for 1h at room temperature and washed three times in TBST, followed by detection by enhanced chemiluminescence (ThermoFisher).

### RNAseq

Total RNA was harvested from 2-3 million HIV-dreGFP infected Jurkat cells exposed to EPZ-719 (500nM) or control (DMSO) using a RNEasy kit (Qiagen). RNA quantity and quality were then analyzed by nanodrop and Tapestation (Agilent) analysis respectively. RNA integrity number (RIN) scores were between 9.9 and 10.0. RNAseq libraries were then prepared using the KAPA Total RNA library prep kit, followed by 50bp paired end sequencing on a Nextseq2000 P2 flow cell. 70-90 million reads were obtained per sample. To analyze the data, the raw reads were first filtered by FastQC and CutAdapt to remove low quality reads. RNAStar was then used to align the reads to Hg38 (patch 17), followed by featurecounts to quantify transcripts. Differentially expressed genes were then identified by DESeq2 [72].

### Quantitative PCR

HIV Gag unspliced RNA and Beta-actin RNA were quantified as previously described [37]. Briefly, cells were washed in PBS then lysed in RLTplus buffer (Qiagen) with 1% beta-mercaptoethanol. RNA was then isolated using an RNEasy plus kit (Qiagen) and eluted into nuclease free water. RNA samples were quantified by nanodrop, then 100ng of RNA was reverse transcribed and amplified using Fastvirus (Thermo, Waltham, MA) and primer sets for HIV Gag RNA (GAG-F: ATCAAGCAGCCATGCAAATGTT, GAG-R: CTGAAGGGTACTAGTAGTTCCTGCTATGTC, GAG-Probe: FAM/ZEN-ACCATCAATGAGGAAGCTGCAGAATGGGA-IBFQ) and Beta-actin (BAC-F: TCACCCACACTGTGCCCATCTACGA, BAC-R: CAGCGGAACCGCTCATTGCCAATGG, BAC-Probe: HEX-ATGCCCTCCCCCATGCCATCCTGCGT-IBFQ). The reaction plate was run on a QS3 (Applied Biosystems, Foster City, CA) real time thermocycler with a 5 minute reverse transcription step at 50°C, followed by 40 cycles of 94°C (3 sec.), 60°C (30 sec.).

### Splicing quantification

Quantification of HIV-1 transcripts was done similarly as described in [47], but adapted to use a random 14 base cDNA primer coupled with Illumina/MiSeq platform sequences. Briefly, the random reverse primer serves as a unique molecular identifier, and also primes across all viral RNAs as well as all cellular RNAs. HIV-1 specific RNAs are polymerase chain reaction (PCR) amplified using a primer just upstream of the major HIV-1 splice donor D1, which also adds Illumina platform sequences, and a downstream primer complementary to a common sequence at the 5’ end of the random reverse primers. The resulting libraries were sequenced using Illumina paired-end 300 base reads. Sequencing data was analyzed using an in-house script (available from the Swanstrom lab, UNC Chapel Hill) that combines information from forward and reverse reads to identify and quantify transcripts that either remain unspliced at D1 or splice to specific acceptors. For reads of sufficient length, transcripts that are spliced or unspliced at D4 (completely or incompletely spliced) can also be quantified.

### Integration site analysis

Genomic DNA was isolated using the DNeasy Blood and Tissue kit (Qiagen) per manufacturers protocols. Integration sites were determined using a linear amplification mediated PCR protocol adapted from methodology originally described in [40]. Duplicate PCR reactions gDNA were subject to linear amplification using the biotinylated primer am948. Resulting linear amplification products were pooled and purified using the NucleoSpin Gel and PCR Purification kit (Takara) with modified protocols for ssDNA (1:2 NTI buffer dilution). All recovered material was subject to streptavidin immunoprecipitation using the Dynabeads kilobaseBINDER Kit (ThermoFisher) using 5uL beads per manufacturers protocol to recover the biotinylated linear amplification product, washed, and followed by second strand synthesis on bead using klenow (New England Biolabs, NEB) as described [40]. Post all enzymatic reactions, DNA-bound beads were washed twice in a tween wash buffer (5mM Tris pH8, 1M NaCl, 0.5mM EDTA, 0.05% Tween-20) and once in 10mM Tris pH8. Double stranded on-bead DNA was digested with a mix of 3 blunt cutting enzymes (SspI-HF, StuI, and HincII) (NEB) to generate blunt-end DNA. A blunt-ended double stranded adapter based of sequences from [40] was generated by annealing two oligos, a sense strand with 5’ unpaired region and antisense with a 3’ blocked end. The adapter was ligated on to digested DNA using the NEB Quick Ligation Kit. Material was subject to on-bead PCR using primers am954/956 using the Platinum SuperFi II Mastermix (ThermoFisher/Life Technologies) for 15 cycles. 2uL of the first round PCR was then subject to nested PCR using am950/955 for 30 cycles using SuperFi II. PCR products were cleaned up using the Takara Nucleospin kit per manufacturers protocol and concentration and size range assessed using the D5000 Screentape (Agilent). 200fmol of each sample was barcoded and subject to library prep using the Native Barcoding Ligation Kit with V14 chemistry (SQK-NBD-114-24) per manufactures instructions. 20fmol of the pooled, barcoded library was loaded onto an MIN114 flow cell and run on the Nanopore Mk1C. Resulting fast5 files were basecalled and demultiplexed using the high accuracy basecalling model for Guppy (Oxford Nanopore). Custom adapters sequences ligated during the protocol were trimmed using Cutadapt [73] (https://cutadapt.readthedocs.io/en/stable/) and successful trimming confirmed using Fastqc (https://www.bioinformatics.babraham.ac.uk/projects/fastqc/) with a custom adapter list pre and post trimming. Resulting reads were initially aligned to the reference HIV genome using minimap2 and non-aligning reads discarded. Reads were then realigned to a hybrid HIV-hg38 genome using minimap2 with the -Y option set to map chimeric reads. Reads were filtered using SAMtools [74] to extract reads with SA tags, sorted, and overlapping reads collapsed using BEDTools merge. Read depth of collapsed features was merged into the BED file using SAMtools bedcov with -d 10 and -c options. Finally, features were overlapped with a reference BED file created using the hg.knowncanonical track from the UCSC Genome Browser using BEDTools intersect and filtered to produce a list of genomic regions with a minimum of 10 nanopore reads. All primers and oligos were synthesized by IDTDNA and are as follows: am948 - /5Biosg/CTTCTAGCCTCYGCTAGTCAAA; am954 - YTC AGC AAG CCG AGT CYT G; am956 - GAC CCG GGA GAT CTG AAT TC AG; am950 - TTT CAR GTC CCT GTT CGG G; am955 - AGT GGC ACA GCA GTT AGG; adapter S - GAC CCG GGA GAT CTG AAT TCA GTG GCA CAG CAG TTA GG; adapter AS - /5Phos/ CCT AAC TGC TGT GCC A/3C6/.

### RNA methylation analysis

To measure the total abundance of m6A RNA in cells, 300ng of total RNA was analyzed using an m6A RNA methylation quantification kit (Epigentek) according to the manufacturer’s protocol. This kit uses a colorimetric enzyme linked immunosorbent assay (ELISA) to detect m6A. ELISA plates were read at 450nm using a SpectraMax M3 plate reader (Molecular Devices). Samples were run in triplicate and compared to a standard curve of m6A. Abundance was calculated as a percentage of total RNA by mass. To measure m6A modification of HIV RNA, an EpiQuikTM CUT&RUN m6A RNA Enrichment (MeRIP) Kit (Epigentek) was used with an input of 10ug of total RNA from each condition and an m6A specific antibody or control IgG. Eluted RNAs were then analyzed using a quantitative PCR assay for an m6A rich region of the HIV RNA within the Env/Rev coding sequence. qPCR was run using an Applied Biosystems QS3. Primer sets used were Forward: GCC CGA AGG AAT AGA AGA AGA A, Reverse: GAT CGT CCC AGA TAA GTG CTA AG, Probe: 56-FAM/TG GAG AGA G/ZEN/A GAC AGA GAC AGA TCC A/3IABkFQ.

## Acknowledgements

This work was supported by the following grants from the National institutes of Health: NIAID #5- R01AI143381, NIAID #5- UM1AI164567, U54-AI15047.

## Supplementary data

**Figure S1:**
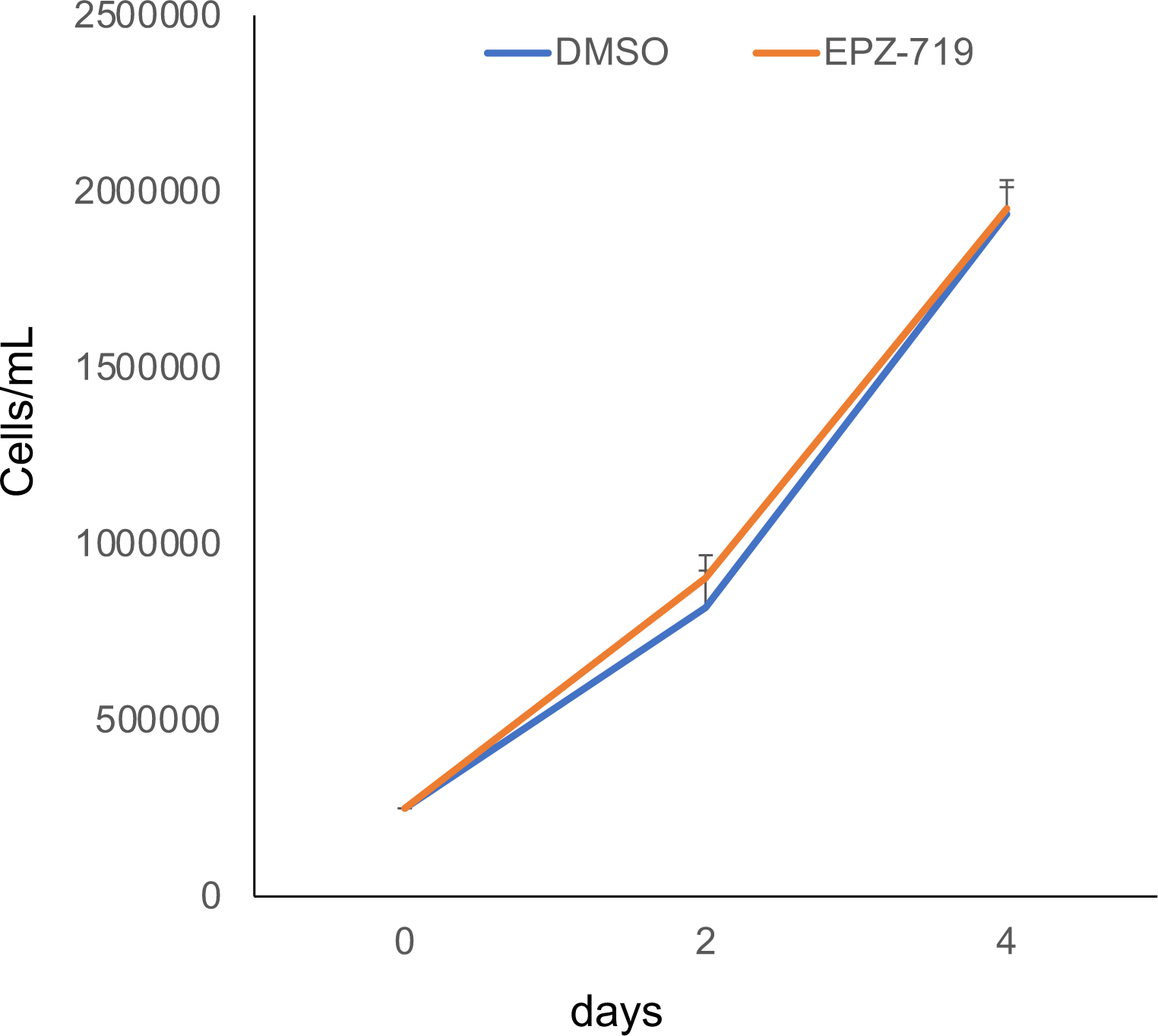
EPZ-719 does not inhibit Jurkat cell growth. Jurkat cells were seeded in RPMI at 250,000 cells per mL and incubate in the presence of 500nM EPZ-719 or DMSO for 4 days. At times indicated, cell density was measured. Each datapoint represents the average of triplicate wells. Error bars represent the standard deviation of the mean.

**Figure S2:**
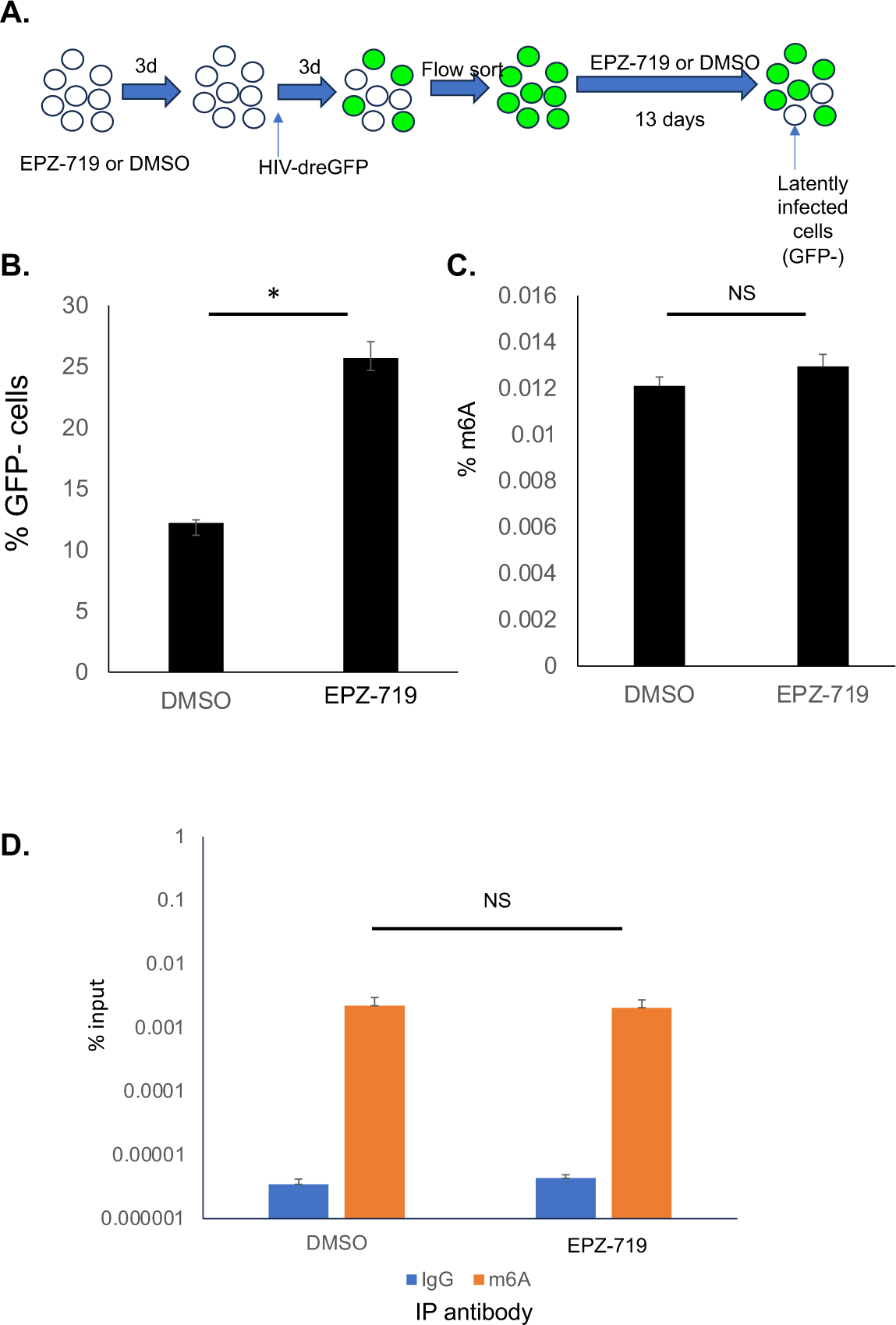
EPZ-719 does not affect global or HIV specific m6A modification of RNA. **A.** Schematic overview of experimental design. EPZ-719 was used at 500nM. **B.** The abundance of latently infected (GFP-) Jurkat cells in the culture was measured by flow cytometry. **C.** The abundance of m6A RNA within the total RNA from DMSO or EPZ-719 exposed was measured by plate-based enzyme linked immunosorbent assay (ELISA). **D**. m6A modification of HIV RNA was examined. Cellular RNA was immunoprecipitated with an m6A-specific antibody or control IgG followed by quantitative RT-PCR for a region of HIV located within the Env/Rev region. Each bar represents the average of biological triplicates. Error bars represent the standard deviation of the mean. NS= not significant, P>0.05 T Test.

